# Cleavage of protein kinase c δ by caspase-3 mediates pro-inflammatory cytokine-induced apoptosis in pancreatic islets

**DOI:** 10.1101/2022.07.20.500512

**Authors:** Jillian Collins, Robert A. Piscopio, Mary E. Reyland, Chelsea G. Johansen, Richard K. P. Benninger, Nikki L. Farnsworth

## Abstract

In type 1 diabetes (T1D), autoreactive immune cells infiltrate the pancreas and secrete pro-inflammatory cytokines that initiate cell death in insulin producing islet β-cells. Protein kinase C δ (PKCδ) plays a role in mediating cytokine-induced β-cell death; however, the exact mechanisms are not well understood. To address this, we utilized an inducible β-cell specific PKCδ KO mouse as well as a small peptide specific inhibitor of PKCδ. We identified a role for PKCδ in mediating cytokine-induced β-cell death and have shown that inhibiting PKCδ protects pancreatic β-cells from cytokine-induced apoptosis in both mouse and human islets. We determined that cytokines induced nuclear translocation and activity of PKCδ and that caspase-3 cleavage of PKCδ may be required for cytokine-mediated islet apoptosis. Further, cytokine-activated PKCδ increases activity both of pro-apoptotic Bax with acute treatment and JNK with prolonged treatment. Overall, our results suggest that PKCδ mediates cytokine-induced apoptosis via nuclear translocation, cleavage by caspase-3, and upregulation of pro-apoptotic signaling in pancreatic β-cells. Combined with the protective effects of PKCδ inhibition with δV1-1, the results of this study will aid in the development of novel therapies to prevent or delay β-cell death and preserve β-cell function in T1D.

## Introduction

Type 1 diabetes (T1D) is characterized by the selective immune-mediated destruction of insulin producing β-cells in pancreatic islets [1, 2]. Clinical onset of T1D results in life-long dependence on exogenous insulin and is associated with long-term complications including, ophthalmic, kidney, cardiovascular, and neurological disorders [3]. While current therapeutic strategies aim to control glucose levels, they do not stop disease progression nor prevent disease complications. Furthermore, current clinical trials to prevent or reverse disease outcomes have had limited success, including modest disease remission and incomplete recovery of normoglycemia [4, 5]. The exact mechanisms regulating β-cell death during T1D pathogenesis are not fully understood, making it difficult to develop effective therapies to preserve endogenous β-cells.

In T1D, pro-inflammatory cytokines secreted from autoreactive immune cells play a crucial role in β-cell apoptosis [6]. Studies have shown that the pro-inflammatory cytokines tumor necrosis factor-α (TNF-α), interleukin 1-beta (IL-1β), and interferon-γ (IFN-γ) work synergistically to induce islet dysfunction and β-cell death *in vitro* [7-9]. In the β-cell, cytokine-mediated apoptosis has been shown to occur through either activation of mitogen-activated protein kinase (MAPK), or activation of nuclear factor kappa B or activation of signal transducer and activator of transcription 1 (STAT1), all of which lead to activation of caspase-3 and apoptosis [6, 10]. Our lab has previously shown that pro-inflammatory cytokines also disrupt gap junction coupling, calcium (Ca^2+^) signaling and impair insulin secretion in both mouse and human islets via upregulation of protein kinase C delta (PKCδ) activity [7].

PKCδ regulates many cellular functions including cell proliferation, differentiation, and apoptosis (11). In response to apoptotic stimuli, such as DNA damaging H_2_O_2_ and IR radiation, tyrosine sites on PKCδ are phosphorylated leading to a conformational change in PKCδ into its active state where it can initiate apoptosis through phosphorylation of downstream pro-apoptotic signaling messengers [11-14]. PKCδ has been shown to mediate fatty acid–induced apoptosis in human β-cells and studies utilizing a mouse model with overexpression of a kinase-negative PKCδ protected against high fat diet mediated apoptosis [15, 16]. Similarly, a whole-body knockout of PKCδ protected against β-cell death in streptozotocin (STZ) induced diabetes in mice *in vivo* and cytokine treated β-cells *in vitro* [17]. However, PKCδ is ubiquitously expressed throughout the body and a global knockout leads to spontaneous Β-cell proliferation resulting in increased autoimmunity [18].

While PKCδ has previously been implicated in mediating β-cell apoptosis in mouse islets, the mechanism by which PKCδ regulates cytokine-induced apoptosis has not been well defined in the β-cell. In normal epithelial cells and some cancer cells, etoposide activated PKCδ is cleaved by caspase-3 into a 40kDa catalytically active fragment that translocates to the nucleus and mediates cell apoptosis [11, 19]. As caspase-3 activation plays a role in cytokine-mediated β-cell death, the goal of this study is to understand the role of caspase-3 in mediating PKCδ regulation of cytokine-induced β-cell death. We hypothesized that cytokines secreted by autoreactive immune cells mediate β-cell death through a signaling pathway that requires activation of PKCδ cleavage by caspase-3 and nuclear accumulation of PKCδ. We utilized isolated islets from a novel β-cell-specific PKCδ knockout mouse or human islets treated with a cell permeable PKCδ specific inhibitor, δV1-1. Identification of novel regulators of cytokine-mediated human β-cell death will provide new avenues towards developing targeted therapies to preserve β-cell mass in T1D.

## Results

### Characterization of the PKCδ^fl/fl^ x MIP-Cre^ER^ mouse

To assess the role of PKCδ in mediating cytokine-induced β-cell death, we developed a β-cell specific knockout of PKCδ (PKCδ-βKO). β-cell specificity was achieved using mouse insulin promoter (MIP) Cre-ER on the C57Bl/6 mouse background, which is expected to produce mosaic expression in β-cells based on previous results with the inducible Cre-ER system [20]. We also bred our MIP-CreER line with a mouse line with tdTomato reporter of Cre recombination inserted into Rosa26 locus as a marker of Cre-induced knockout [21]. Islets isolated from PKCδ-βKO mice 2 weeks after tamoxifen injection showed robust expression of tdTomato in 76.5±5% of cells, where tdTomato is a reporter for Cre recombination and knockout of PKCδ (Figure 1A). For reference, mouse islets are composed of an average of 60-80% β-cells [22]. PKCδ-βKO islets had 58% less PKCδ protein compared to control (PKCδ^fl/fl^) as determined by western blot (p=0.015, Figure 1B and C). No significant alterations in glucose tolerance levels were observed between the PKCδ-KO and control mice using an intraperitoneal glucose tolerance test (Figure 1D).

**Figure 1.**
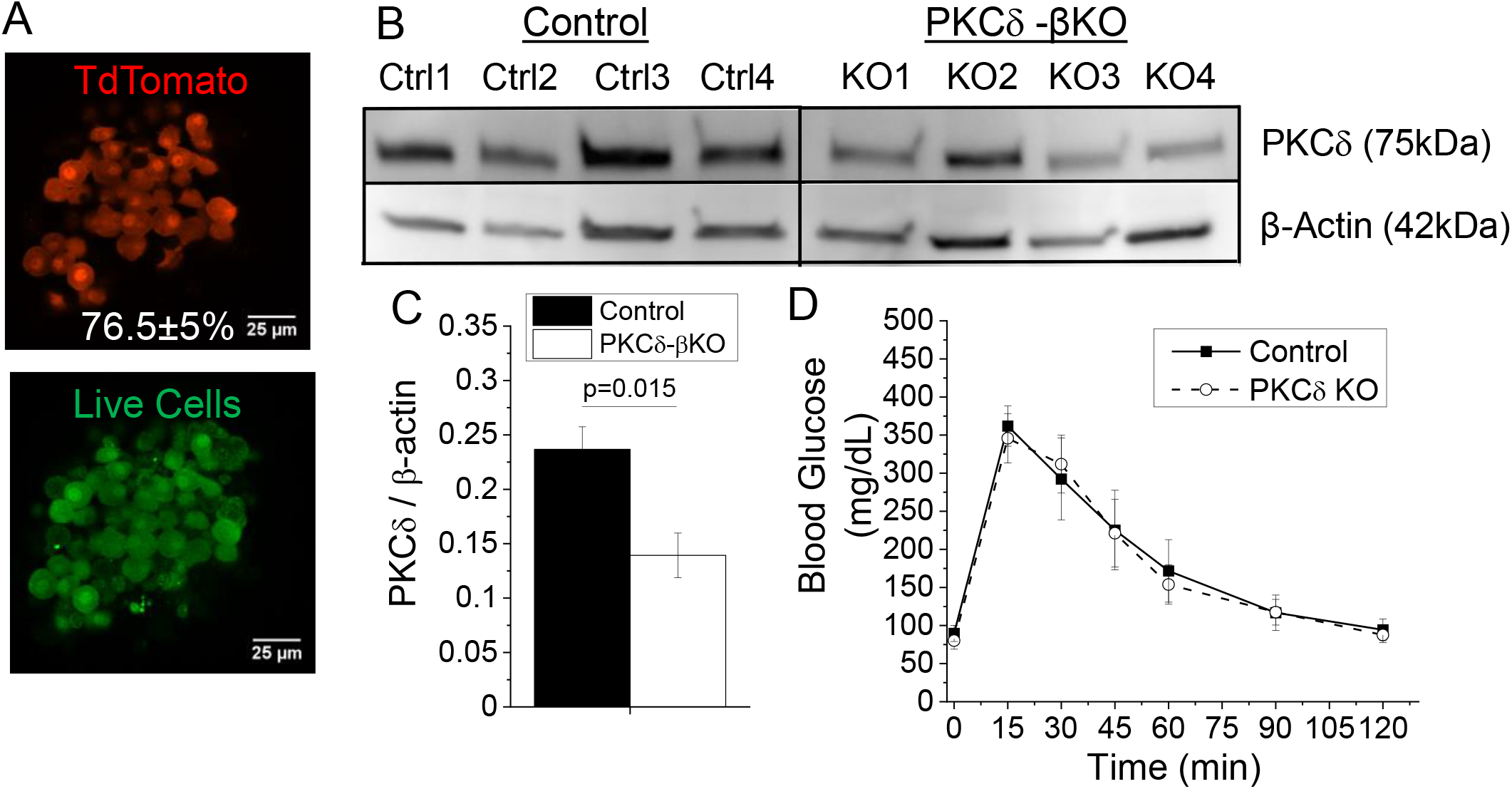
β-cell specific inducible KO of PKC8 (PKC8-βKO). (A) Representative images of tdTomato fluorescence (red) in PKC**8**-βKO islets stained with a live cell dye (green) 2 weeks after tamoxifen injection, where 76.5±5% of all cells were tdTomato positive (n=4). (B) Representative western blot of PKC**8** and β-actin controls from islets isolated from 4 control and 4 age and gender matched PKC**8**-βKO mice 2 weeks after tamoxifen injection. Images are from the same blot that has been cropped to remove the empty lane between samples. (C) Quantification of western blots from panel A of PKC**8** protein normalized to β-actin in islet isolated from control and PKC**8**-βKO mice two weeks after tamoxifen injection (n=4). P<0.05 is statistically significant based on a paired-sample t-test. (D) Blood glucose levels during a glucose tolerance test in 8-15 week old control and age and gender matched control (MIPCre^ER^ negative) and PKC**8**-βKO mice 2 weeks after tamoxifen injection following a glucose bolus (n=3).

### PKCδ mediates cytokine-induced apoptosis in mouse and human islets

To determine if PKCδ is required for cytokine-induced apoptosis in the β-cell, isolated PKCδ-βKO, MIP Cre-ER positive, and control islets were treated with or without a cocktail of proinflammatory cytokines TNF-α (10ng/ml), IL-1β (5 ng/ml), and IFN-γ (100ng/ml) for 24hr. A significant increase in apoptotic cells, as measured by the apoptosis specific dye YO-PRO1 (Figure 2A), was observed in cytokine treated control and MIP Cre-ER positive islets compared to untreated control islets (p<0.001, Figure 2B). The percentage of apoptotic cells was significantly reduced in the cytokine treated PKCδ-βKO islets compared to MIP Cre-ER negative control or MIP Cre-ER positive islets (p=0.003, Figure 2B). Islets were also treated with a zinc^2+^ sensor for quantifying insulin expressing cells in addition to NucBlue Live cell stain and propidium iodide for dead cells (Figure 2C). The percentage of insulin positive dead cells was significantly less in the IL-1β, TNF-α, and IFNγ cytokine cocktail treated PKCδ-βKO islets compared to control islets (p=0.04, Figure 2D). Mouse C57Bl/6 and human islets were also treated with or without cytokine cocktail and with or without the PKCδ peptide inhibitor δV1-1. Both mouse (p<0.001) and human (p=0.02) islets treated with cytokine cocktail displayed a decrease in the percentage of apoptotic cells with δV1-1 treatment compared to cytokine only control (Figure 2E and F). This data supports a role for PKCδ in mediating cytokine-induced apoptosis, specifically in pancreatic β-cells.

**Figure 2.**
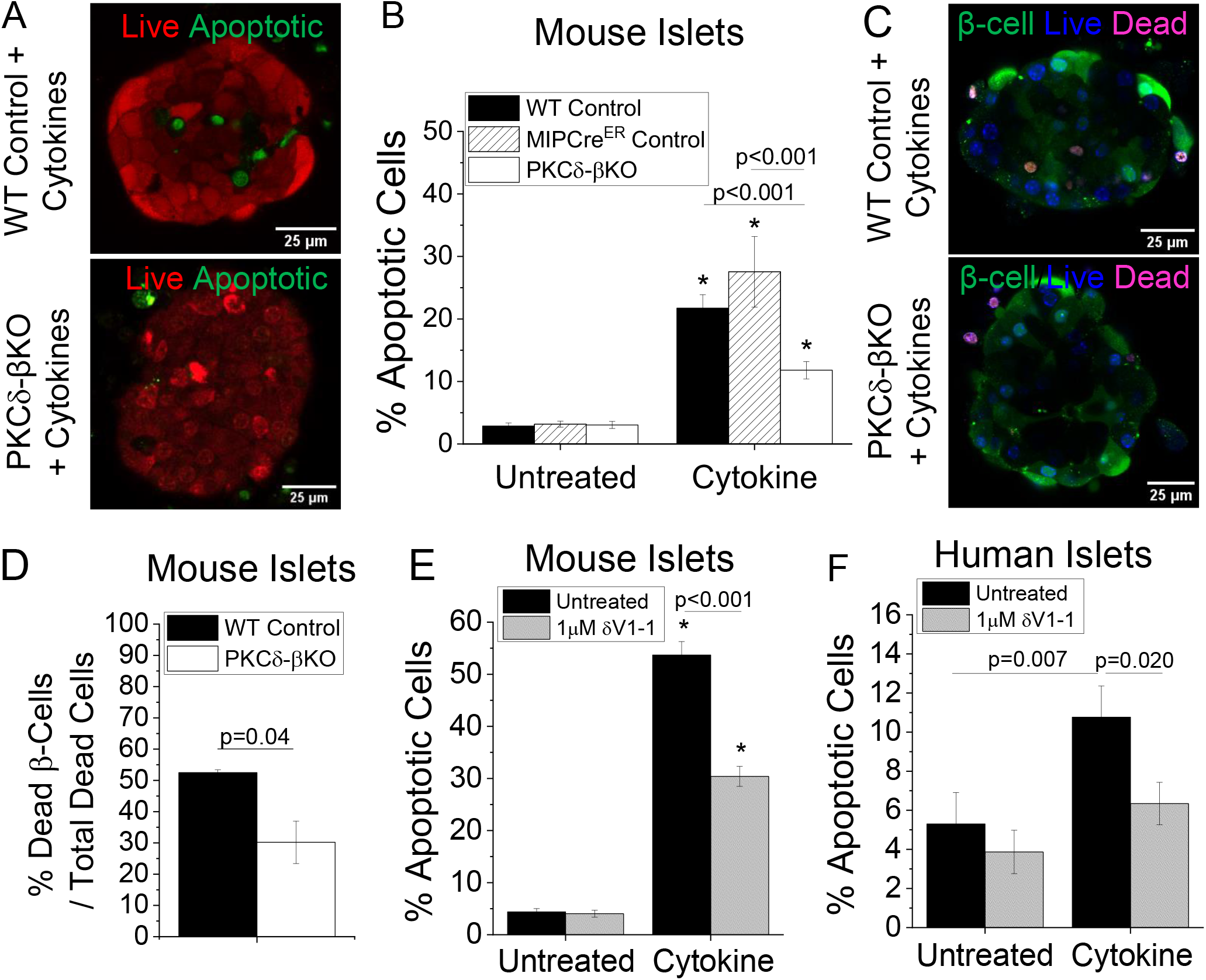
PKCδ mediates cytokine-induced islet apoptosis. (A) Representative images of live (red) and apoptotic (green) cells as measured by NucRed Live and YO-PRO1 respectively in WT control and PKC8-βKO islets treated with cytokines for 24h. (B) Percent apoptotic cells in isolated control, MIP Cre-ER positive or PKC8-βKO mouse islets treated for 24h with a cytokine cocktail compared to untreated islets in 4-5 islets per experiment (n=16). (C) Representative images of FluoZin-3 staining of β-cells (green), live cells labeled with NucBlue live (blue), and dead cells labeled with propidium iodide (PI, red). (D) Percent insulin positive apoptotic cells in isolated control or PKC8-βKO mouse islets treated for 24h with a cytokine cocktail compared to untreated islets (n=3). (E) Percent apoptotic cells in C57Bl/6 mouse islets treated for 24h with a cytokine cocktail and the PKC8 inhibitor 8V1-1 in 4-5 islets per experiment as indicated (n=7). (F) Percent apoptotic cells in human islets from four donors treated for 24h with a cytokine cocktail and the PKC8 inhibitor 8V1-1 as indicated (n=4). * Indicates a significant difference in cytokine treated samples compared to untreated controls for islets from each respective mouse. p<0.05 indicates statistical significance based on ANOVA with Tukey’s post-hoc analysis.

### Cytokine Treatment Induces Nuclear Translocation and Activity of PKCδ in Islets

Nuclear translocation of PKCδ is an indication of activation and is associated with the pro-apoptotic functions of PKCδ [23]. To investigate whether a cytokine cocktail induced nuclear translocation of PKCδ in pancreatic islet β-cells, PKCδ-βKO islets were transduced with an adenovirus to overexpress GFP fused to the c-terminus of PKCδ (GFP-PKCδ, Supplemental Figure 1) or with an adenovirus to express GFP alone as a control overnight and transduced cells were cultured with or without cytokine cocktail for 24 hours [23]. GFP-PKCδ was expressed in PKCδ-βKO islets to avoid cell death due to supra-physiological levels of PKCδ. Prior to imaging, islets were stained with NucRed live to determine nuclear versus cytosolic PKCδ localization. In transduced untreated islets we observed diffuse GFP-PKCδ signal throughout the cytosol and the nucleus, with nuclear GFP intensity generally lower than cytosolic GFP (Figure 3A). We observed an increase in GFP fluorescence in the cell nuclei with cytokine cocktail treatment compared to untreated controls (Figure 3A, red arrows). Furthermore, at both 3hr (p=0.034) and 24hr (p<0.001) timepoints, the fluorescence intensity in the nucleus normalized to the cytosol significantly increased in cytokine treated islets compared to untreated controls (Figure 3B). We also transfected cells with GFP-only using an adenoviral vector to confirm that GFP was not influencing the observed nuclear translocation with cytokine cocktail treatment. No significant change in the fluorescence intensity in the nucleus normalized to the cytosol was observed in PKCδ-βKO islets treated with GFP-only virus with cytokine cocktail treatment compared to untreated controls (Figure 3C).

**Figure 3.**
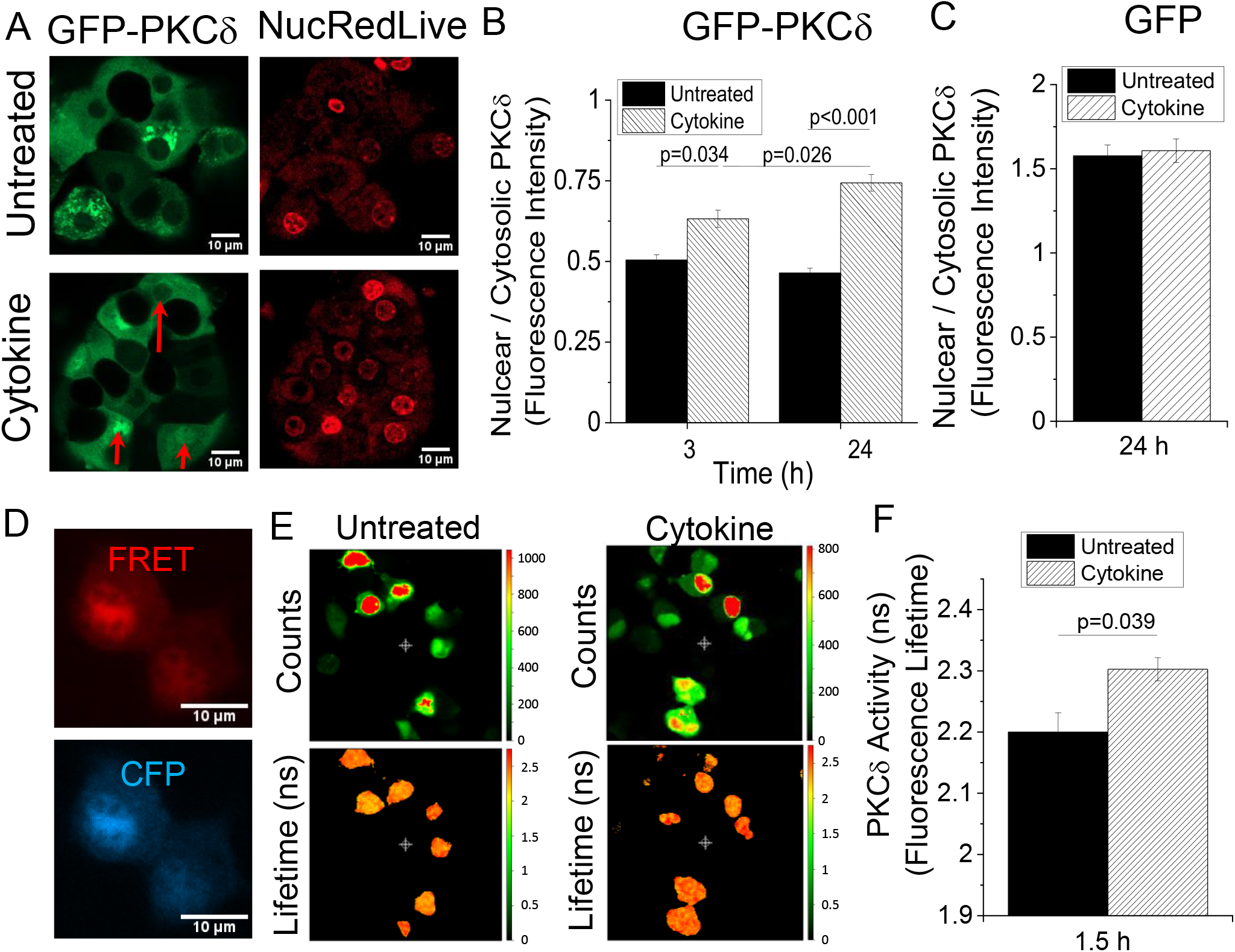
Cytokines induce nuclear translocation and activity of PKCδ. (A) Representative images of GFP-PKCδ (green) in PKCδ-βKO islets co-stained with NucRedLive (red) and either untreated or treated with cytokines for 24 h. Red arrows point to nuclei with increased fluorescence intensity compared to cytoplasm in cytokine treated samples. (B) Quantification of GFP intensity in the nucleus normalized to intensity in the cytoplasm of each cell in 4-5 islets per experiment (3h n=4, 24h n=10). (C) Quantification of GFP intensity in PKCδ-βKO islets co-stained with NucRedLive (red) and transduced with GFP only and treated with cytokines for 24 hours (n=4). (D) Representative images YFP and CFP emission with CFP excitation using the FRET sensor for PKCδ (δCKAR) targeted to the nucleus. (E) Representative images of fluorescence lifetime imaging in MIN6 cells transfected with the FRET-based PKCδ activity sensor in E. Images on the top represent the number of photon counts and images on the bottom represent the calculated fluorescence lifetime in ns for untreated (left panel) and cytokine treated (right panel) MIN6 cells after 1.5h culture. (F) Quantification of PKCδ activity by fluorescence lifetime imaging of the FRET sensor for PKCδ activity in the nucleus in MIN6 cells either untreated or treated with cytokines for 1.5 hours (n=3). In panels B and C, p<0.05 indicates a significant difference as determined by ANOVA with Tukey’s post-hoc analysis. In panel E, p<0.05 indicates a significant difference as determined by students T-test.

We next measured changes in nuclear activity of PKCδ with cytokine cocktail treatment using a nuclear targeted PKCδ specific FRET-based activity sensor (δCKAR) in MIN6 cells. Expression of δCKAR was primarily localized to the nucleus (Figure 3D). The average lifetime in the nuclear region of the cells (Figure 3E) was measured using fluorescence lifetime imaging (FLIM) using a threshold on photon counts to calculate to determine changes in FRET with cytokine cocktail treatment. For reference, when no FRET is occurring and PKCδ is maximally active the expected lifetime is ∼2.9ns; however, when FRET occurs with 100% efficiency and there is no PKCδ activity the anticipated lifetime is ∼1.9ns [24]. We found that the fluorescence lifetime of the δCKAR sensor increased in cells treated with cytokine cocktail, which indicates an increase in PKCδ activity in the nuclei of cytokine cocktail treated cells compared to untreated controls after 1.5 hours of treatment (p=0.039, Figure 3F).

### Cytokine-Induced Cleavage of PKCδ by Caspase-3

In several cell types, cleavage of PKCδ yields a constitutively active fragment that localizes to the nucleus to promote apoptosis [25]. Therefore, to determine if caspase-3 cleaves PKCδ during cytokine cocktail-induced apoptosis, and if caspase cleavage is necessary for cytokine-induced apoptosis, PKCδ-βKO islets were transduced with an adenovirus encoding a GFP-PKCδ peptide, where the sequence that can be cleaved by caspase-3 has been mutated to prevent PKCδ cleavage (CM-GFP-PKCδ). Islets treated with viruses coding for GFP-only, GFP-PKCδ, or CM-GFP-PKCδ were cultured for 24h with or without cytokine cocktail (Supplemental Figure 1). Cytokine cocktail caused significant increases in apoptosis under all conditions tested (Figure 4A). No significant differences were observed in cytokine cocktail-induced apoptosis levels between the PKCδ-βKO alone and GFP-only treatment (Figure 4A), indicating that GFP does not contribute to islet apoptosis. There was a significant increase in cytokine cocktail-induced apoptosis in islets expressing GFP-PKCδ compared to islets with no virus pre-treatment (p<0.001, Figure 4A), indicating normal function of the GFP-PKCδ peptide in the islet. The cytokine cocktail-treated CM-GFP-PKCδ islets showed a significant reduction in the percentage of dead cells compared to cytokine cocktail treated GFP-PKCδ islets (p<0.001, Figure 4A) and similar levels to GFP-only treated cells indicating the inhibition of caspase-3 cleavage can mitigate PKCδ mediated apoptosis. Translocation of the CM-GFP-PKCδ to the nucleus was also measured as previously described and an increase in the nuclear to cytosolic ratio of the GFP intensities was observed with cytokine cocktail treatment (p=0.009, Figure 4B), indicating increased nuclear translocation of the intact PKCδ determined as described above. Additionally, western blot confirmed that similar levels of GFP-PKCδ and CM-GFP-PKCδ expression were obtained in PKCδ-βKO islets (Figure 4C and 4D). These results show that caspase cleavage of PKCδ is required for PKCδ-mediated apoptosis; however, cleavage is not required for nuclear translocation with cytokine cocktail treatment.

**Figure 4.**
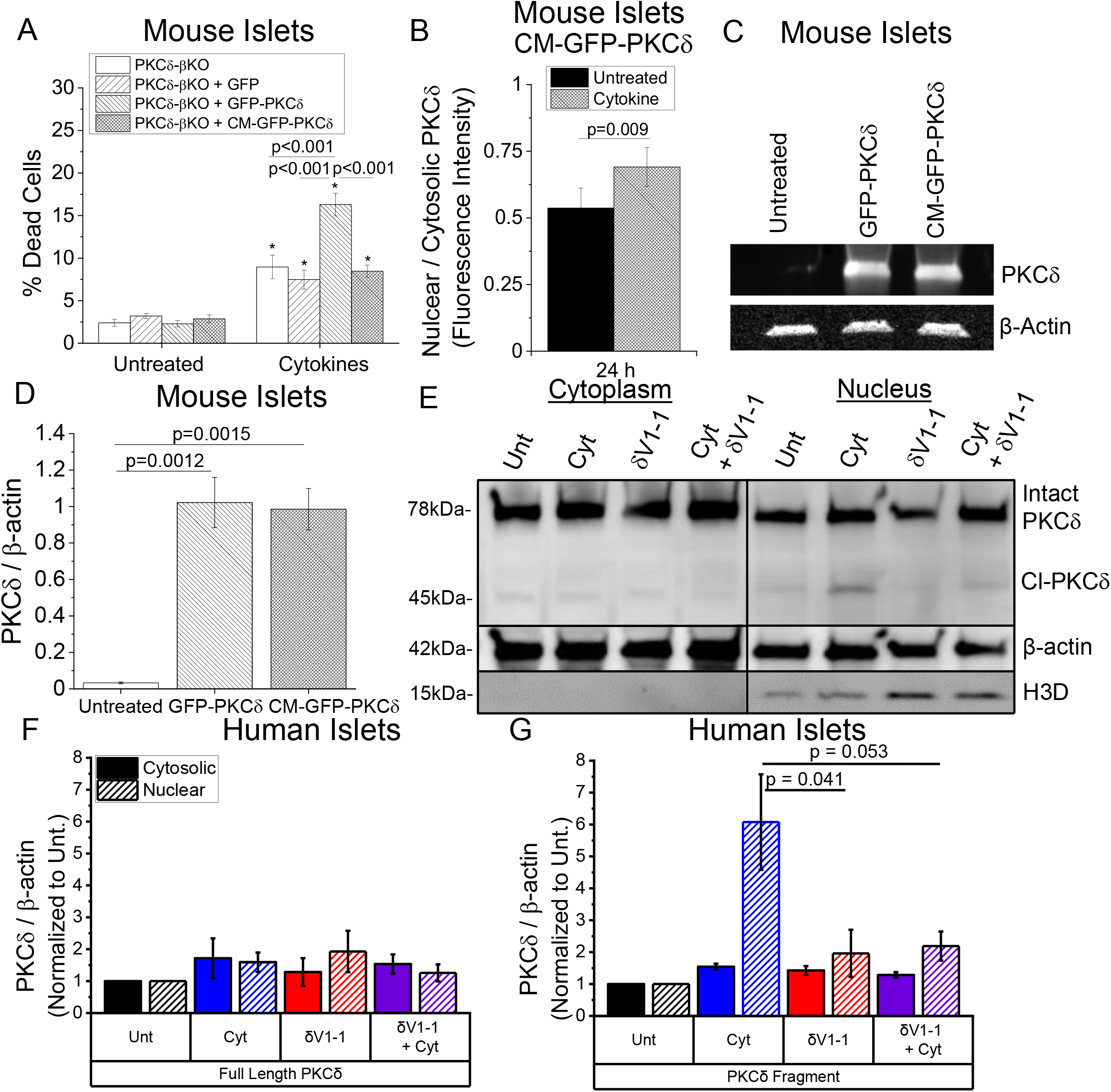
Cleavage of PKCδ is required for cytokine-mediated islet apoptosis. (A) Percent apoptotic cells in mouse PKCδ-βKO islets treated for 24h with a cytokine cocktail and either GFP-only, GFP-PKCδ, or CM-GFP-PKCδ virus compared to untreated islets in 4-5 islets per experiment (n=4). * Indicates a significant difference from untreated islets (p<0.01). (B) Quantification of GFP fluorescence intensity in the nucleus normalized to intensity in the cytosol in PKCδ-βKO mouse islets transfected with the CM-GFP-PKCδ virus as in A and either untreated or treated with cytokines for 24h in 4-5 islets per experiment (n=3). (C) Representative western blot of GFP-PKCδ, CM-GFP-PKCδ, and β-actin expression in PKCδ-βKO islets compared to control islets not treated with virus 24 h after treatment. (D) Quantification of the blots in E of GFP-PKC or CM-GFP-PKCδ in PKCδ-βKO islets normalized to β-actin expression (n=3). (E) Representative western blot of PKCδ, β-actin, and nuclear marker histone cluster 1 H3D in lysates from human islets treated for 24 hours with or without cytokines and with or without the PKCδ inhibitor δV1-1. Islet lysates were fractionated into cytosolic and nuclear fractions. Images are from the same blot that has been cropped to place the cytoplasmic samples on the left. Two bands were identified for PKCδ staining, one at 75kDa representing full length PKCδ and one at 45kDa representing a cleaved version of PKCδ (Cl-PKC8). Quantification of the blots in A of either intact PKCδ (75kDa) (F) or the 45kDa PKCδ fragment (G) in both the cytosolic and nuclear fractions normalized to β-actin and further normalized to untreated control samples. p<0.05 indicates a significant difference based on ANOVA.

To determine if caspase-3 play a similar role in mediating cytokine-induced apoptosis and PKCδ cleavage in human islets, subcellular fractionation of lysates and western blot were used to observe changes in both intact (78kDa) and cleaved fragments (Cl-PKCδ, 45kDa) of PKCδ in both the cytosolic and nuclear fractionations of human islet lysates treated with or without cytokine cocktail and with or without the PKCδ inhibitor δV1-1 (Figure 4E). Staining of histone cluster 1 H3D in the nuclear fraction and not in the cytosolic fraction confirmed separation of the cytosolic and nuclear lysate (Figure 4E). Quantification of the blot stained for PKCδ and β-actin revealed no significant changes in full length PKCδ (78kDa) in the cytosolic fraction or the nuclear fraction under any treatment, when normalized to untreated samples (Figure 4F). This suggests that PKCδ translocation measured in Figure 3A and B was mainly the PKCδ fragment, as GFP is fused to the c-terminus (Supplemental Figure 1) and will identify both intact and cleaved PKCδ. While a fragment (45kDa) of PKCδ was observed in the cytosol for all samples (Figure 4E), no significant changes were observed in the PKCδ fragment in the cytosol under any treatment, when normalized to untreated samples (Figure 4G). A significant increase in the PKCδ fragment was observed in the nuclear fraction of cytokine cocktail treated islets compared to untreated controls (based on 95% confidence interval, Figure 4G). Treatment with the PKCδ inhibitor δV1-1 reduced cytokine-mediated increases in the PKCδ fragment in the nucleus (p=0.053, Figure 4G), supporting a reduction in PKCδ cleavage. These results support that δV1-1 inhibits activation of PKCδ as previous studies have shown that the caspase-3 cleavage site on PKCδ is only accessible after conversion of PKCδ from its immature folded conformation to an unfolded and activated conformation [14, 25-27]. Overall, these results support that cytokine-mediated cleavage of PKCδ is required for β-cell death in both mouse and human islets.

To further verify the role of caspase-3 in cytokine-mediated apoptosis and regulation of PKCδ, mouse and human islets were treated for 24hr with the caspase-3 inhibitor ZEVD-FMK with or without cytokine cocktail (Figure 5). Inhibiting caspase-3 protected against cytokine-induced apoptosis in both mouse (p<0.001) and human islets (Figure 5A and B). To determine if inhibiting caspase-3 altered cleavage of PKCδ with and without cytokine treatment, western blot analysis of PKCδ and β-actin was performed on control and PKCδ-βKO mouse islet lysates treated for 24 hours with or without ZEVD-FMK and with or without cytokines (Figure 5C). While inhibiting caspase-3 had no significant impact on the level of full length PKCδ normalized to β-actin (Figure 5D), we observed a decrease in the level of the PKCδ fragment (45kDa) normalized to β-actin (Figure 5E) or normalized to intact PKCδ (Figure 5F) in ZEVD-FMK treated islets compared to untreated controls. A similar decrease in levels of the PKCδ fragment was observed in islets treated with cytokines and ZEVD-FMK compared to cytokine cocktail alone (p=0.05, Figure 5E). Overall, these results support the conclusion that caspase-3 cleavage of PKCδ is required for PKCδ mediation of cytokine-induced β-cell apoptosis in both mouse and human islets.

**Figure 5.**
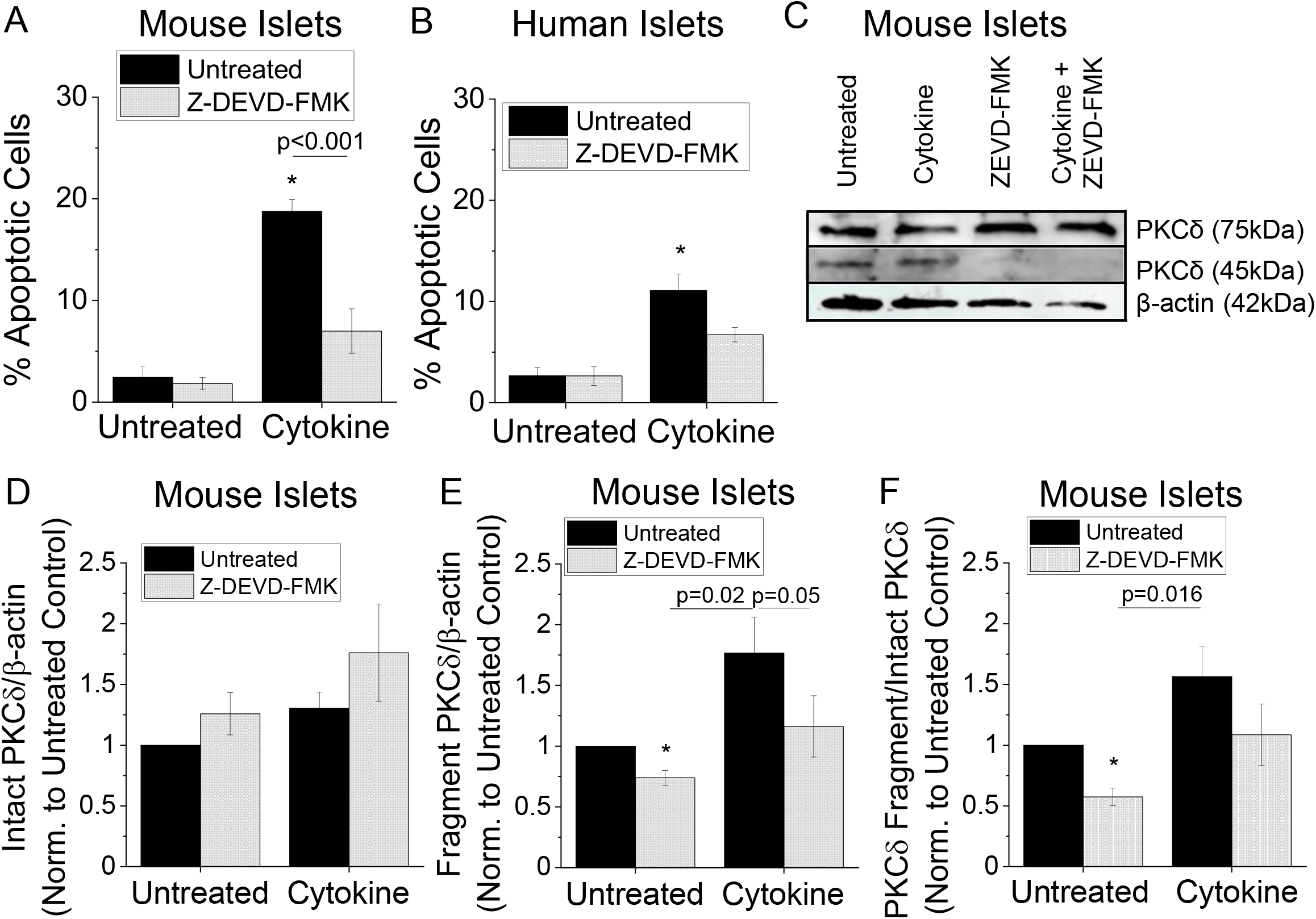
Inhibiting caspase-3 protects against cytokine induced apoptosis and prevents PKCδ cleavage. (A) Percent apoptotic cells in mouse C57Bl/6 islets treated for 24h with a cytokine cocktail and the caspase-3 inhibitor Z-DEVD-FMK as indicated compared to untreated islets (n=4). (B) Percent apoptotic cells in human islets treated for 24h with a cytokine cocktail and the caspase-3 inhibitor Z-DEVD-FMK as indicated compared to untreated islets (n=3). * Indicates a significant difference from untreated islets (p<0.01). (C) Representative western blots of PKCδ and β-actin in lysates from C57Bl/6 mouse islets treated with or without cytokines and with or without the caspase-3 inhibitor Z-DEVD-FMK for 24 hours as in A and B. (D) Quantification of the blots in C of intact (75kDa) PKCδ normalized to β-actin, where data was normalized to untreated controls for each experiment (n=5). (E) Quantification of the blots in C of the PKCδ fragment (45kDa) normalized to β-actin, where data was normalized to untreated controls for each experiment (n=5). (F) Quantification of the blots in C of the PKCδ fragment (45kDa) normalized to intact PKCδ (75kDa), where data was normalized to untreated controls for each experiment (n=5). In D-F, * indicates a significant difference from untreated controls as determine by 95% CI. p<0.05 in all panels indicates a significant difference based on ANOVA.

### Cytokine-Activated PKCδ Mediates Pro-Apoptotic Signaling

Finally, we investigated PKCδ-mediated changes in pro-apoptotic signaling, including Bax and C-Jun N-terminal kinases (JNKs), with acute (1h) and prolonged (24h) exposure to cytokine cocktail. Control and PKCδ-βKO islets were either untreated or treated with cytokine cocktail for 1 or 24 hours and levels of Bax, phospho-JNK (p-JNK), JNK, and β-actin were analyzed via western blot (Figure 6). After 1 hour cytokine cocktail treatment, we found slight increases in Bax levels in WT islets treated with cytokine cocktail compared to untreated controls (Figure 6A and C). The cytokine-mediated increase in Bax was abolished in PKCδ-βKO islets compared to WT controls (p=0.018, Figure 6A and C). JNK1 (46kDa) and JNK2 (54kDa) were both observed with JNK staining of the western blot (Figure 6B). We found that after 1h cytokine treatment, no significant changes in p-JNK normalized to total JNK1 (46kDa) were observed between cytokine cocktail treated islets and untreated controls or between WT and PKCδ-βKO islets (Figure 6B and D). After 24 hours of cytokine cocktail treatment, we found that Bax levels were similar in both control and PKCδ-βKO islets either untreated or treated with cytokine cocktail for 24hr (Figure 6E and G). An increase in p-JNK/JNK1 was observed with cytokine cocktail treatment in both the control and PKCδ-βKO islets (p=0.016, Figure 6F and H). Only the quantification of JNK1 is shown as JNK2 levels did not significantly change in control versus PKCδ-βKO islets either in untreated or cytokine cocktail treated conditions (data not shown). Overall, this data supports a role for PKCδ in differentially regulating cytokine-mediated pro-apoptotic signaling at 1 hour versus 24 hours in the islet.

**Figure 6.**
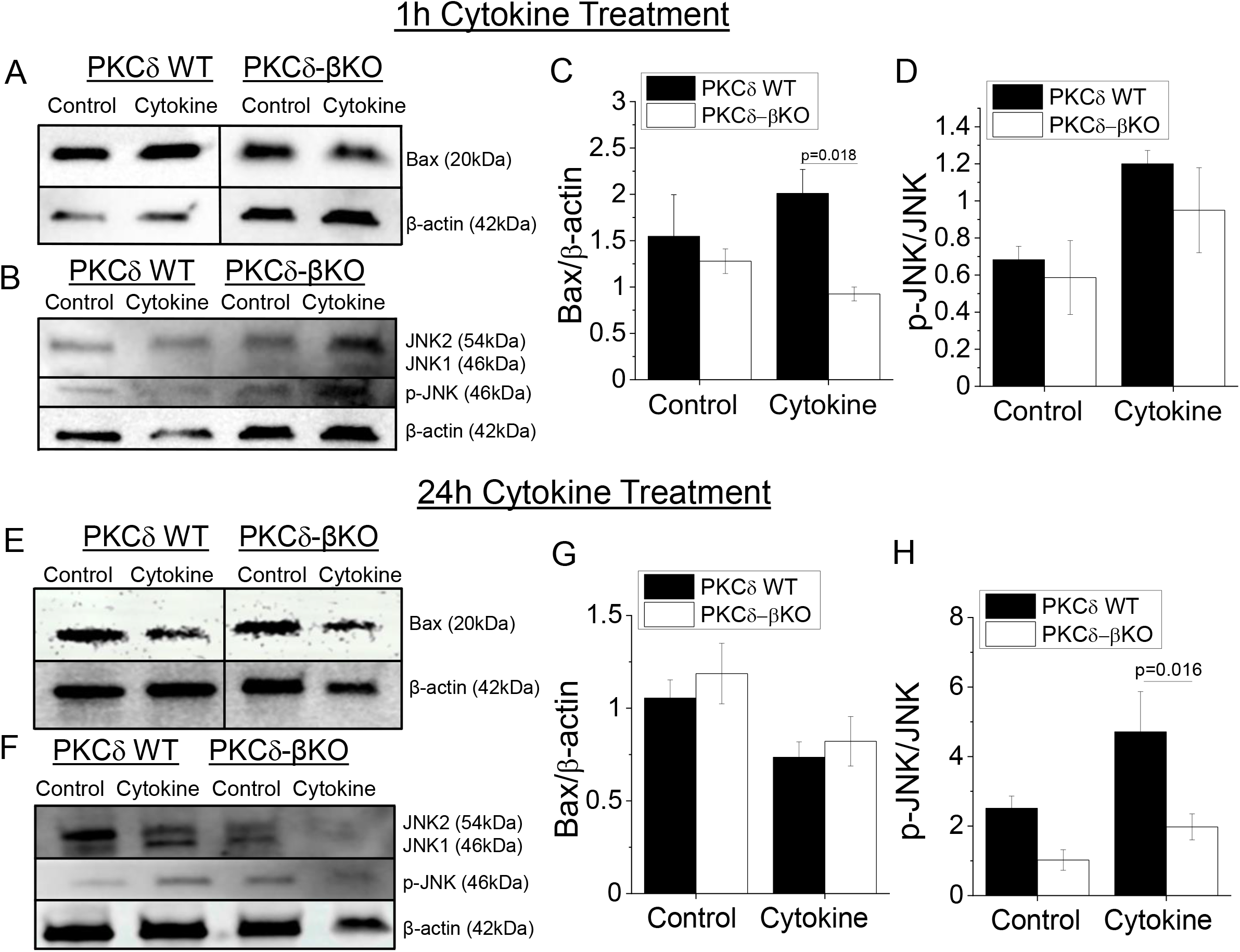
PKCδ mediates pro-apoptotic signaling. (A) Representative western blots of Bax and β-actin in lysates from control and PKCδ-βKO mouse islets treated with or without cytokines for 1 hour. Blots were cropped to put images of WT islet lysates to the left of PKCδ-βKO islet lysates to match the other panels. (B) Representative western blot of JNK, p-JNK, and β-actin in lysates from control and PKCδ-βKO mouse islets treated with or without cytokines for 1 hour. (C) Quantification of the blots in A, where Bax levels were normalized to β-actin for each condition (n=9-12). (D) Quantification of the blots in B, where p-JNK was normalized to JNK1 (46kDa) levels (n=5-7). (E) Representative western blots of Bax and β-actin in lysates from control and PKCδ-βKO mouse islets treated with or without cytokines for 24 hours. Blots were cropped to put images of WT islet lysates to the left of PKCδ-βKO islet lysates to match the other panels. (F) Representative western blot of JNK, p-JNK, and β-actin in lysates from control and PKCδ-βKO mouse islets treated with or without cytokines for 24 hours. (G) Quantification of the blots in E, where Bax levels were normalized to β-actin for each condition (n=4). (H) Quantification of the blots in F, where p-JNK was normalized to JNK1 (46kDa) levels (n=4). In panels C, D, G and H p<0.05 indicates a significant difference based on ANOVA with Tukey’s post-hoc analysis.

## Discussion

The goal of this study was to elucidate the mechanism by which PKCδ mediates cytokine-induced apoptosis and identify a role for caspase-3 cleavage of PKCδ in regulating cytokine-mediated β-cell death in pancreatic islets. Utilizing an inducible β-cell specific PKCδ KO mouse as well as a small peptide specific inhibitor of PKCδ, we confirmed that PKCδ activity mediates cytokine-induced islet apoptosis in both mouse and human islets. We determined that cytokines induce nuclear translocation and activity of PKCδ. Further, our results support that caspase-3 cleavage of PKCδ is required for cytokine-mediated islet apoptosis. Ultimately, cytokine-activated PKCδ was shown to regulate pro-apoptotic Bax expression with acute cytokine treatment and JNK activity with prolonged cytokine treatment, suggesting a downstream mechanism of PKCδ-mediated apoptosis as shown in Figure 7. Combined with the protective effects of PKCδ inhibition with δV1-1, the results of this study will aid in the development of novel therapies to prevent or delay β-cell death and preserve β-cell function in T1D.

**Figure 7.**
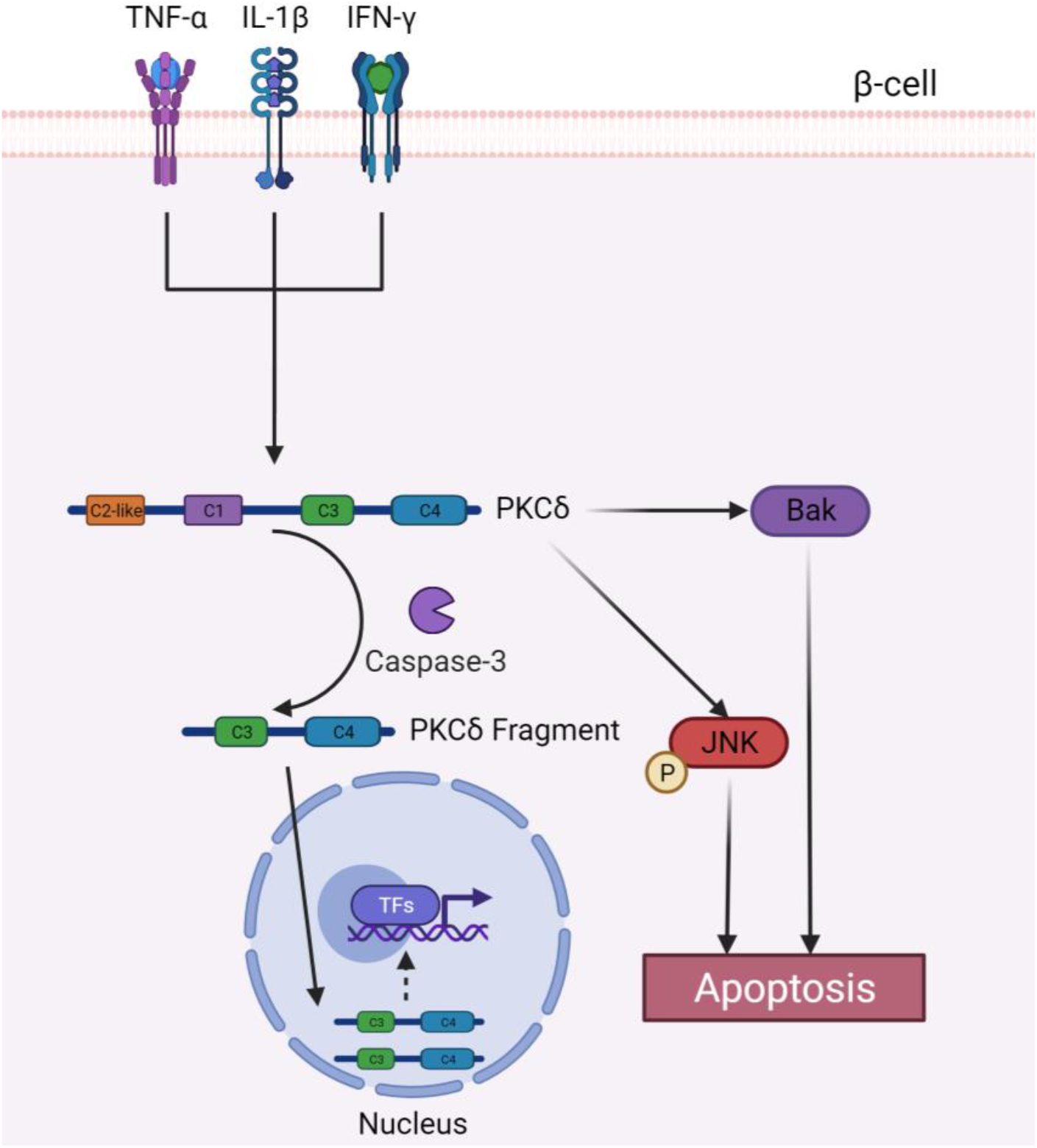
Proposed mechanism for PKCδ regulation of cytokine-induced β-cell death. Cytokine activated PKCδ is cleaved by caspase-3 into a 45kDa fragment that translocates to and accumulates in the nucleus. Cytokine activated PKCδ promotes apoptosis through activation of Bax and JNK. We propose that catalytic activity of PKCδ in the cytosol activates Bax while nuclear PKCδ activity upregulates expression of kinases that activate JNK downstream of PKCδ.

### Inhibition of PKCδ Protects Against Cytokine-Induced Death

Pro-inflammatory cytokines play a critical role in mediating islet cell death and dysfunction in T1D [6, 7, 28]. Several studies have identified a potential role for PKCδ in mediating cytokine-induced β-cell death. For example, a knockout of PKCδ, or expression of a kinase negative PKCδ, protected against cytokine-induced islet death and improved glucose homeostasis and β-cell function in high-fat diet mice *in vitro* [16, 17, 29]. While these findings support a role for PKCδ in mediating cytokine-induced islet death, the results are confounded by the effects from a whole body PKCδ KO as PKCδ is ubiquitously expressed [18]. To determine the role of PKCδ specifically in mediating β-cell death, we generated and characterized a novel mouse line with an inducible β-cell specific KO of PKCδ (PKCδ-βKO). Contrary to previously published results indicating that loss of PKCδ impairs glucose tolerance, our PKCδ-βKO mice were normoglycemic compared to control controls [30, 31]. While this may be due incomplete loss of PKCδ protein in the islets of our transgenic mouse, this may also be indicative of confounding effects of a whole-body KO from loss of PKCδ in the liver, where previous studies have indicated that global PKCδ KO impairs insulin sensitivity [32, 33]. Our data supports Cre-mediated KO of PKCδ in ∼76% of islet cells, which is in agreement with previous studies indicating that 60-80% of rodent islets are composed of β-cells [22]. It has been previously reported that PKCδ is present in α-cells, which comprise up to 20% of a pancreatic islet, that may also explain the incomplete knockout of PKCδ in whole islet lysates [34]. Overall, our results support knockout of PKCδ in the majority of β-cells that will allow us to probe the role of PKCδ in mediating cytokine-induced apoptosis.

Our results indicate that KO of PKCδ significantly protects against cytokine-induced apoptosis in mouse islets. To specifically analyze β-cell apoptosis, we used the YO-PRO1 dye, which specifically labels early apoptotic cells through selective permeability and has been previously shown to overlap with TUNEL staining of apoptotic cells and not propidium iodide staining of membrane integrity, supporting specific protection against β-cell apoptosis [35]. Furthermore, this protection was specific to β-cells, as β-cell specific cytokine-induced apoptosis was reduced in PKCδ-βKO islets compared to WT controls, demonstrating the efficacy of our β-cell specific knockout mouse.

To confirm our results with a genetic KO of PKCδ, we also utilized a cell permeable specific inhibitor of PKCδ, δV1-1, an FDA approved peptide that specifically inhibits PKCδ binding to membrane associated RACK anchor proteins that facilitate PKCδ activation by inducing a conformation change [36]. Our results in human islets treated with cytokines and with or without δV1-1 show a reduction in PKCδ cleavage and translocation to the nucleus. The caspase cleavage site and nuclear localization sequence of PKCδ are only accessible after PKCδ has been activated by phosphorylation and undergone a conformational change; therefore, our results support inhibition of cytokine-mediated PKCδ activation by δV1-1 [23, 37]. We found that inhibiting PKCδ activation with δV1-1 protects against cytokine-induced islet apoptosis in both mouse and human islets. Collectively, this strongly supports a role for PKCδ in mediating cytokine-induced apoptosis specifically in the β-cell and our results in human islets suggest that cytokine-induced activity of PKCδ may promote β-cell death under inflammatory conditions associated with diabetes [6]. Future studies are warranted to investigate the role of PKCδ in β-cell death during diabetes pathogenesis and to determine therapeutic efficacy of δV1-1 in protecting against β-cell death in T1D.

### Cleavage and Nuclear Translocation of PKCδ is Required for Cytokine-Mediated β-Cell Apoptosis

Full length PKCδ is an isoenzyme of ∼78kDa and is comprised of a regulatory and catalytic domain separated by a hinge region [14, 38]. In the presence of apoptotic stimuli, such as ionizing radiation and etoposide, PKCδ has been shown to be proteolytically cleaved into a constitutively active catalytic fragment (∼45kDa) which can undergo translocation from the cytosol to the nucleus via a nuclear localization sequence [23, 26, 27]. In this study nuclear translocation and accumulation of PKCδ was observed with cytokine-treated islets via expression of a fusion protein with GFP tagged to the C-terminus, showing localization of both full length and cleaved PKCδ (see Supplemental Figure 1). Furthermore, we demonstrated that PKCδ was cleaved into a smaller 45kDa fragment in the presence of pro-inflammatory cytokines that accumulate in the nucleus. While we were unable to distinguish between the activity of full length PKCδ and the 45kDa fragment in the nucleus, we observed cytokine-mediated increases in nuclear PKCδ activity that correlates with increases accumulation of the 45kDa fragment. Our results indicate that caspase-3 cleaves cytokine-activated PKCδ into the observed 45kDa fragment and that cleavage is required for PKCδ mediated cytokine-induced apoptosis in the islet. While were unable to determine if PKCδ is cleaved before or after nuclear translocation, DeVries-Seimon et al. provide evidence that supports intact PKCδ translocates to the nucleus before being cleaved by caspase-3 in etoposide treated parC5 cells [19]. Caspase-3 has also been shown to undergo nuclear translocation in α-Fas or etoposide treated HepG2 cells only after activation, further suggesting that cleavage of PKCδ occurs in the nucleus [39]. This is supported by our results where expression of a mutant PKCδ that cannot be cleaved by caspase-3 translocates to the nucleus in the presence of cytokines but translocation of the full length PKCδ did not increase β-cell apoptosis. Overall, our results suggest that PKCδ mediates cytokine-induced β-cell apoptosis via nuclear translocation and caspase-3 cleavage into a 45kDa fragment. This novel mechanism of cytokine-induced apoptosis involving PKCδ and caspase-3 may present a unique target to prevent inflammation-induced β-cell death in diabetes.

Furthermore, our results support an increase in PKCδ activity in the nucleus, suggesting that PKCδ may mediate cytokine-induced apoptosis through phosphorylation and subsequent activation of downstream mediators of apoptosis either directly or through activation of transcription factors that regulate pro-apoptotic genes. Previous studies in keratinocytes have shown that PKCδ mediates activation of the STAT3 transcription factor, which is regulated by cytokine signaling and increase expression of pro-apoptotic genes [40]. In the β-cell, PKCδ has also been shown to activate the FOXO1 transcription factor in response to free fatty acids, leading to β-cell apoptosis [16]. Taken together, our results supported further investigation into the role PKCδ in regulating cytokine-induced apoptosis via upregulation of pro-apoptotic signaling pathways.

### Cytokine-Activated PKCδ Regulates Pro-apoptotic Signaling

PKCδ regulation of pro-apoptotic signaling has been characterized in several cell types. For example, in lung cancer cells exposed to ionized radiation, activation of PKCδ leads to downstream activation of pro-apoptotic Bax, ultimately resulting in apoptosis [41]. Additionally, jun N-terminal kinases (JNKs) are a protein kinase family that regulate cell death, where cytokine-induced inflammation has been shown to specifically activate JNK1 [42, 43]. This led us to investigate a potential role for PKCδ in mediating the activation of Bax and JNK with cytokine treatment. Our results are consistent with previous studies in human islets that showed increases in Bax activity between 1-3 hours and increases in p-JNK at 1h and 24h after treatment with IL-1β and IFN-γ [44]. Overall, our results in PKCδ-βKO islets support a role for activated PKCδ in regulating apoptotic signaling via a fast mechanism that activates Bax after 1h of cytokine treatment and via a slower mechanism that activates JNK1 after 24h of cytokine treatment. In human keratinocytes, activation of the catalytic subunit of PKCδ by 4-hydroxytamoxifen leads to redistribution and activation of Bax that initiates downstream cytochrome-c release from the mitochondrial and cell apoptosis [45]. Furthermore, previous studies in cancer cells have shown that activated PKCδ up-regulates expression of apoptosis signal-regulating kinase 1 (ASK1), which directly activates JNK via phosphorylation [46]. Taken all together with our results, this supports a role for PKCδ in regulating cytokine-induced apoptosis via acute activation of Bax through catalytic activity in the cytosol and longer term upregulation of genes encoding for kinases that activate JNK. Future studies will further investigate the mechanisms by which PKCδ activates Bax and JNK as well as potential regulation of anti-apoptotic signaling. Further characterization of the downstream signaling pathways mediated by PKCδ will provide novel insight into cellular targets to protect pancreatic islets from cytokine induced death in T1D.

## Conclusion

To summarize, we identified a role for PKCδ in mediating cytokine-induced β-cell death and have shown that inhibiting PKCδ protects pancreatic β-cells from cytokine-induced apoptosis in both mouse and human islets. Pro-inflammatory cytokine treatment results in PKCδ translocation from the cytosol to the nucleus were PKCδ is cleaved by caspase-3. Cleavage by caspase-3 was also found to be required for PKCδ mediated cytokine-induced cell death. Furthermore, our data suggests that cytokine-activated PKCδ increases the activity of the pro-apoptotic Bax with acute cytokine treatment and JNK1 with prolonged cytokine treatment. Overall, the results from this study provide novel insight into the role of PKCδ in mediating cytokine-induced apoptosis in pancreatic islets and supports a role for PKCδ in mediating cytokine-induced apoptosis in T1D in humans. Furthermore, the results of this study have clinical implications as targeted inhibition of this novel PKCδ mediated apoptotic pathway may preserve β-cell mass in T1D.

### Experimental procedures

#### Mice

Animals were housed in a temperature-controlled facility with access to food and water ad libitum and were on a 12h light/dark cycle. PKCδ^LoxP/LoxP^ (PKCδ^fl/fl^) animals were a generous gift from Dr. Ronald Kahn [32]. Mouse insulin promoter (MIP) Cre^Er^ [47], Rosa26 tdTomato [21], and C57Bl/6 mice were purchased from The Jackson Laboratory (Bar Harbour, ME). To generate a β-cell specific knockout of PKCδ, PKCδ^fl/fl^ and MIP-Cre^Er^ mice were bred, which we will refer to as control (PKCδ^fl/fl^) and PKCδ-βKO (PKCδ^fl/fl^MIPCre^Er^). control and PKCδ-βKO were bred in house and genotyping was performed by Transnetyx (Cordova, TN) using real-time PCR with the following primer sets: Cre (Forward: TTAATCCATATTGGCAGAACGAAAAC, Reverse: CAGGCTAAGTGCCTTCTCTACA), PKCδ^fl/fl^ (Forward: CTCCCCACCGATTAGTGTTGAAAA, Reverse: GGACGTAAACTCCTCTTCAGACCTA), PKCδ^WT^ (Forward: GGCAGTTATCTGACTCTTGCAGCT, Reverse: CCAAATAGGAACAACAGGTGCCTCT).

#### Human Islets

Human islets were obtained from the Integrated Islet Distribution Program (IIDP) from the following donors:

| Donor ID | Viability | Purity | Age | Sex | BMI | Race |
| --- | --- | --- | --- | --- | --- | --- |
| SAMN17928660 | 90% | 96% | 62y | M | 25.9 | White |
| SAMN18196260 | 95% | 95% | 41y | F | 28.0 | Black |
| SAMN18092805 | 90% | 96% | 56y | M | 26.0 | Asian |
| SAMN23079315 | 90% | 90% | 37y | F | 30.2 | White |

#### Cell Lines

MIN6 cells were cultured in DMEM with 10% FBS and 5% penicillin/streptomycin at 37°C and were passaged at ∼80% confluency using 0.25% trypsin. MIN6 cells used in this study ranged from passage 23-30 as previous studies have shown loss of glucose responsiveness after passage 35 in culture [48].

#### Materials

Recombinant mouse and human pro-inflammatory cytokines (TNF-α (410-MT, 210-TA), IL-1β (401-ML, 201-LB), and IFN-γ (485-MI, 285-IF) were purchased from R&D systems (Minneapolis, MN). RPMI 1640, DMEM, Hanks Balanced Salt Solution (HBSS), fetal bovine serum (FBS), penicillin, streptomycin, YO-PRO1 (Y3603), NucBlue Live ReadyProbes Reagant (R37605), fluorescein diacetate, lipofectamine^TM^ 3000 transfection reagent (L3000015), Halt^TM^ Protease & Phosphatase Inhibitor Cocktail (1861281), FluoZin^TM^-3 (F24195), and caspase inhibitor Z-DEVD-FMK (FMK004) were purchased from Fisher Scientific (Pittsburgh, PA). Collagenase (C9263), tamoxifen, corn oil, D-glucose, ethylenediaminetetraacetic acid (EDTA, EDS), ethylene glycol-bis(2-amino-ethylether)-N,N,N’N’-tetraacetic acid (EGTA), HEPES, Magnesium chloride hexahydrate (H_12_Cl_2_MgO_6_), propidium iodide, were purchased from Millipore-Sigma (Saint Louis, MO). Potassium chloride was purchased from Avantor Performance Materials (Center Valley, PA). PKCδ inhibitor, δV1-1 was purchased from AnaSpec Inc. (AS-65111, Fremont, CA). Western blot buffers and reagents were purchased from Azure biosystems (Dublin, CA). Bacterial vectors encoding FRET sensors for PKCδ activity targeted to the nucleus (δCKAR) were a kind gift from Dr. Alexandra Newton [49, 50].

#### Fasting Glucose Tolerance Test (GTT)

Mice were fasted for 16hr with access to water ad libitum and fasting glucose measurements were taken from the tail vein prior to intraperitoneal injection of 200mg/kg glucose. Blood glucose was monitored over 2hr post glucose bolus via the tail vein.

#### Islet Isolation and Culture

Control and PKCδ-βKO mice were injected with 50mg/kg tamoxifen in corn oil once daily for five consecutive days to induce recombination. Islets were isolated from 8–12-week-old C57Bl/6 or 8–12-week-old age and sex matched control and PKCδ-βKO mice two weeks after the first tamoxifen injection. For islet isolation, animals were injected with 100mg/kg ketamine and 8mg/kg xylazine and euthanized via exsanguination. Islets were isolated by injecting the pancreas with 12.5mg/mL collagenase, pancreas removal, and subsequent enzymatic digestion at 37°C. Islets were handpicked into 1640 RPMI Medium (Millipore-Sigma) with 10% FBS, 10,000 U/mL Penicillin and 10,000μg/mL Streptomycin and incubated overnight at 37°C and 5% CO_2_. Both mouse and human islets were cultured overnight at 37°C and 5% CO_2_ prior to use in experiments.

#### Adenoviral Vectors

Adenoviral vectors for expression of GFP-only, GFP fused to the c-terminus of PKCδ (GFP-PKCδ), and PKCδ with a D→A327 point mutation that can’t be cleaved by caspase-3 with GFP fused to the c-terminus (CM-GFP-PKCδ) were generated by Dr. Mary Reyland as previously described [23, 51] and the expressed proteins are shown in Supplemental Figure 1. Briefly, mouse PKCδ was cloned into the pEGFP-N1 adenoviral vector and the point mutation in PKCδ in the CM-GFP-PKCδ vector was generated by PCR using the primers and site mutagenesis kit descried by DeVries, Neville, and Reyland 2002 [23].

#### Islet Treatments and Apoptosis Measurements

PKCδ-KO and control islets were cultured for 24hr either untreated or treated with a cytokine cocktail at 1X relative cytokine concentration (1 RCC, 10ng/ml TNF-α, 5ng/ml IL-1β, 100ng/ml IFN-γ). Human and C57Bl/6 mouse islets were treated with a cytokine cocktail containing human or mouse recombinant cytokines respectively and 1µM of a PKCδ inhibitor δV1-1 for 24hr. Islets were stained with either propidium iodide or YO-PRO1, a stain that identifies apoptotic cells [35], and NucBlue or fluorescein diacetate (FDA). Imaging was performed either on a Leica STELLARIS 5 LIAchroic laser supply unit, with a 40X water immersion objective. 405nm, 488nm, and 514nm solid state lasers were used for excitation and emission was collected with HyD spectral detectors or on a Zeiss LSM 780 with 488nm or 514nm excitation laser. All islets were imaged as a Z-stack consisting of 3 images 8-10μm apart and live/dead cells were counted manually in ImageJ (NIH). To assess the extent PKCδ was knocked out of the pancreatic β-cells in the PKCδ-KO mice, untreated and cytokine treated islets were stained with FluoZin-3 (1μl/ml) to visualize insulin+ cells. To determine if caspase-3 mediates cytokine-induced apoptosis, islets from female C57Bl/6 mice or isolated human islets were cultured for 24hr with or without cytokines and with or without the caspase-3 inhibitor Z-DEVD-FMK (10μl/ml). Islets were co-stained with YO-PRO1 (1hr) and NucBlue as described above and imaged with a Leica Stellaris confocal microscope with a 40X water immersion objective. Live and dead nuclei were manually counted in ImageJ in 5-10 islets per treatment per experiment. To determine if cleavage of PKCδ by caspase-3 mediates cytokine-induced β-cell death, PKCδ-KO islets were either untreated or treated with a GFP-only expression virus (GFP), GFP-PKCδ, or a GFP-tagged cleavage mutant of PKCδ (CM-GFP-PKCδ, 1:2000) which cannot be cleaved by caspase-3, all for 24hr with or without cytokines. GFP fusion protein expression was confirmed by western blot. Islets were co-stained with YO-PRO1 and NucBlue as described above and imaged with a Leica Stellaris confocal microscope. Live and dead nuclei were manually counted in ImageJ in 5-10 islets per treatment per experiment.

#### PKCδ Translocation to the Nucleus

PKCδ localization and translocation was determined with a GFP adenoviral vector, a GFP-tagged PKCδ adenoviral vector (GFP-PKCδ) or a GFP-tagged cleavage mutant of PKCδ which cannot be cleaved by caspase-3 (CM-GFP-PKCδ) for 24h. Viral vectors were generously provided by Dr. Mary Reyland and were constructed as described above (Supplemental Figure 1). Mouse PKCδ-KO islets were used to reduce potential deleterious contributions from endogenous PKCδ. Isolated islets were cultured with GFP-PKCδ for 24 hours prior to culture for 3 or 24hr with or without cytokines. The GFP-PKCδ virus was also cultured simultaneously with cytokine treatment. Islets were co-stained with NucRed or NucBlue prior to imaging. Islets were imaged either on a Leica Stellaris confocal or a Zeiss LSM 800 confocal with 40X water immersion objective and a 405nm/488nm/514nm excitation laser for each respective dye. Nuclear translocation was determined in ImageJ by normalizing the GFP pixel intensity within each nucleus as determined by colocalization with NucRed or NucBlue, against its respective cytosolic GFP pixel intensity. The ratio of nuclear to cytosolic fluorescence intensity for all GFP-expressing cells was averaged for each islet, in 3-5 islets per experiment.

#### Determining Nuclear PKCδ Activity

PKCδ activity in the nucleus was determined with a PKCδ specific Förster resonance energy transfer (FRET)-based kinase activity sensor (δCKAR) containing a nuclear localization sequence, which was generously given by Dr. Alexandra Newtown and previously described [50]. The δCKAR construct was transfected into MIN6 cells using lipofectamine^TM^ 3000 transfection reagent per the manufacturer’s instructions. Transfected islets were treated for 1.5hr with or without cytokines and changes in FRET were measured with time-resolved fluorescence lifetime imaging (FLIM) on a Zeiss LSM 780 confocal with a tunable infrared Coherent Chameleon Ultra II laser (Zeiss, Oberkochen, Germany. FLIM imaging was done in 2-photon mode. Fluorescence was excited at 720nm with fs-pulses generated by a Coherent Chameleon laser. The microscope is equipped with an ISS FastFLIM acquisition unit and a Becker and Hickl SPC-150E TCSPC acquisition card which allow for fluorescence lifetime imaging in frequency domain and time correlated single photon counting mode respectively in 2 photon imaging. FLIM images were acquired for MIN6 cells over 3 independent experiments. The average FRET-sensor lifetime in 3-10 MIN6 cells was analyzed in ISS VistaVision software with a threshold of 100-200 photon counts applied prior to lifetime calculations with a logarithmic fit and bin size of 1, and the average lifetime was calculated in a manually drawn region of interest for each cell.

#### Subcellular Fractionation

Subcellular fractionation was conducted as previously described on humans islets [7]. Briefly, human islets were washed with 1X TBS and placed in cold fractionation buffer containing 20mM HEPES, 10mM KCl, 2mM MgCl_2_, 1mM EDTA, 1mM EGTA, 1mM DTT and 1mM phosphatase and protease inhibitor cocktail. Samples were incubated on ice for 15min before being passed through a 25G needle 20 times, incubated on ice for 20min and centrifuged at 3,000 rpm for 5min. The supernatant, which contained the cytosolic content was transferred to a separate tube and the nuclear pellet was re-suspended with fractionation buffer, passed through a 25G needle, and centrifuged at 3,000 rpm for 10min. The supernatant was discarded, and the pellet was re-suspended with TBS/0.1wt% SDS and sonicated.

#### Western Blotting

Mouse and human islets were cultured for 1 or 24hr with treatments as indicated. Mouse islets were washed once with PBS and lysed by sonication for 30s in lysis buffer containing 100mM NaCl, 50 mM TrisHCl, 10 mM MgCl2, 1 mM dithiothreitol, and with a protease & phosphatase inhibitor cocktail (5ul/ml). Human islets were fractionated into cytosolic and nuclear components as described above. Protein concentration was measured with Pierce BCA Protein Assay Kit (Thermo Scientific, Rockford, IL) per manufacturer’s instructions. Samples were run on 4–15% mini-PROTEAN® TGX protein gels (Bio-Rad) and transferred to either a PVDF (AC2105; Azure Biosystems) or nitrocellulose (AC2107; Azure Biosystems) membrane. PVDF membranes were blocked in chemi-blot blocking buffer (AC2148; Azure Biosystems) for 2hr and probed with the following antibodies for >2hr at 4°C, all at a 1:1,000 dilution: anti-Bax (ab32503; Abcam, Cambridge, MA), anti-PKCδ (ab182126; Abcam); anti-β-actin (NC9426659; Fisher Scientific); anti-p-JNK (sc-6254, Santa Cruz, CA), and anti-JNK (sc-7345; Santa Cruz). Secondary anti-rabbit (102649-670; VWR) or anti-mouse (626520; Fisher Scientific) horseradish peroxidase-conjugated antibodies diluted 1:5000-1:10,000 for 2hr at room temperature. All membranes were imaged using an Azure c600 imaging system (AC6001; Azure Biosystems) and protein quantification was performed in ImageJ using densitometric analysis. Membranes containing fractionated samples were were stripped and re-stained with nuclear marker histone cluster 1 H3D (sc-134355; Santa Cruz) at a 1:100 dilution and anti-mouse horseradish peroxidase-conjugated secondary diluted 1:10,000. Blot were imaged and quantified as described above.

#### Statistical Analysis

Statistics were performed using Origin software (OriginLabs, Northampton, MA). Two sample t-test and one-way ANOVA with Tukey’s post hoc analysis were performed as indicated. A p-value of <0.05 was considered statistically significant.

## Study Approval

All experiments with mice were approved by the University of Colorado Denver Institutional Animal Care and Use Committee (Protocols 000929 and 00024).

## Data availability

All data presented in this manuscript will be made available upon request to the corresponding author.

## Author Contributions

J.C. conducted experiments, acquired data, analyzed data, and wrote the manuscript. R.A.P. acquired data and analyzed data. M.E.R. designed experiments, assisted in data analysis, and edited the manuscript. C.G.J. acquired and analyzed data. R.K.P.B. designed experiments and edited the manuscript. N.L.F. designed experiments, acquired data, analyzed data, and edited the manuscript.

## Acknowledgements

The authors would like to acknowledge the funding sources that made this work possible, including the following grants: Juvenile Diabetes Research Foundation grants 3-APF-2019-749-A-N and 1-FAC-2020-891-A-N to NLF and 5-CDA-2014-198-A-N to RKPB, Colorado Clinical and Translational Science Institute grant CO-M-19-133 to NLF, American Diabetes Association grant 7-21-JDF-020 to NLF, and National Institute of Diabetes and Digestive and Kidney Diseases grant F32 DK1022706 to NLF and R01 DK102950 and R01 DK106412 to RKPB. We would also like to acknowledge the University of Colorado Diabetes Research Center Islet Isolation Core funded by NIH grant P30-DK116073 and the Advanced Light Microscopy Core facility partly funded by NIH grants P30 NS048154, P30 DK116073. R01DE015648 and R01DE027517 to MER. The content is solely the responsibility of the authors and does not necessarily represent the official views of the National Institutes of Health.

---

**Supplemental Figure 1:**
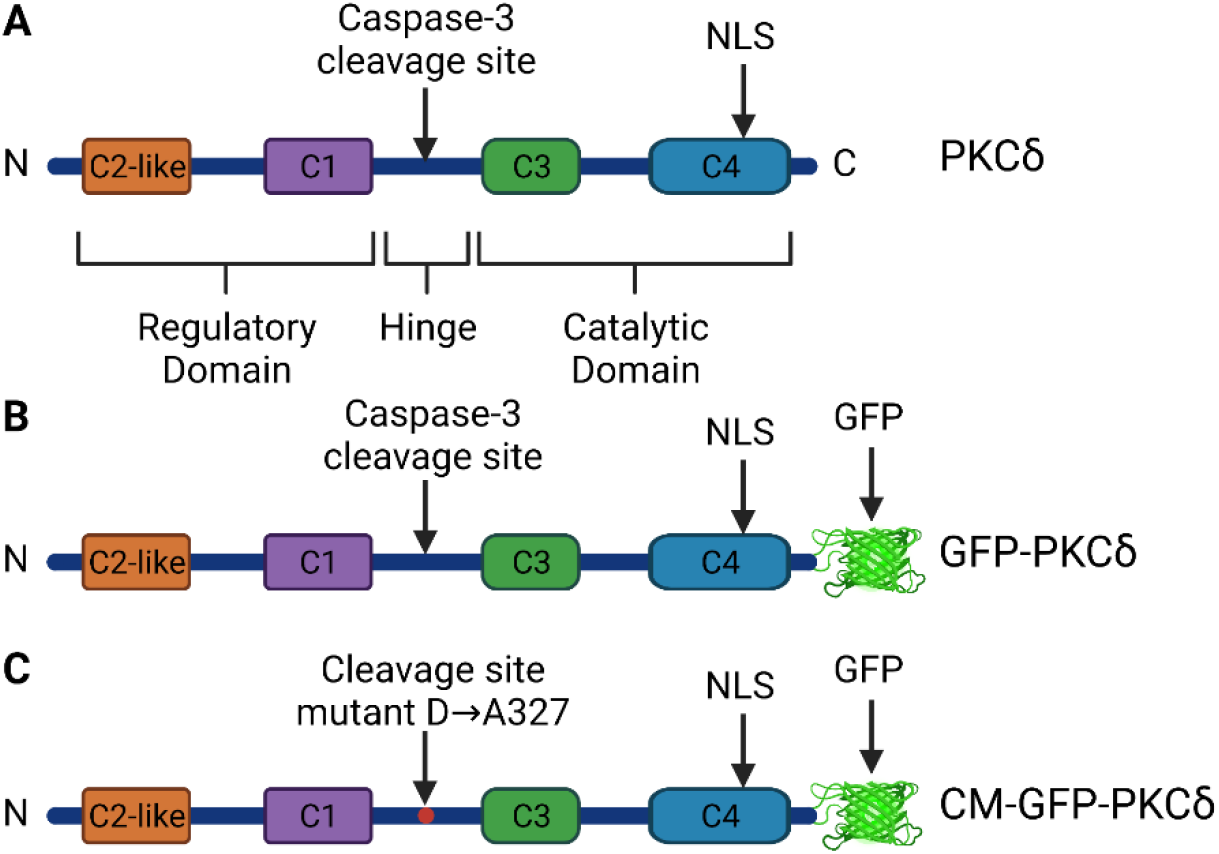
Schematic diagrams of native PKCδ and Modified PKCδ Expressed Through Viral Vectors. (A) Native PKCδ contains regulatory and catalytic domains connected by a hinge domain. The nuclear localization sequence is located in the C4 region of the catalytic domain near the C-terminus and the site where PKCδ can be cleaved by caspase-3 is located in the hinge region. (B) The GFP-PKCδ viral construct expresses PKCδ with EGFP fused to the C-terminus of the protein. (C) The CM-GFP-PKCδ viral construct expresses PKCδ with D327 in the caspase-3 cleavage site mutated to A327 and EGFP fused to the C-terminus.

