## Supplemental Figure 1 for "Cleavage of protein kinase c δ by caspase-3 mediates pro-inflammatory cytokine-induced apoptosis in pancreatic islets"

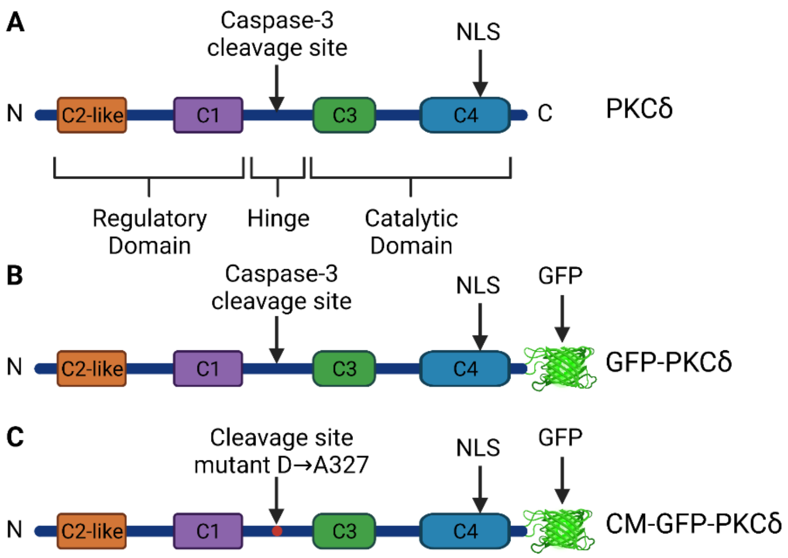


**Supplemental Figure 1: Schematic diagrams of native PKCδ and Modified PKCδ Expressed Through Viral Vectors.** (A) Native PKCδ contains regulatory and catalytic domains connected by a hinge domain. The nuclear localization sequence is located in the C4 region of the catalytic domain near the C-terminus and the site where PKCδ can be cleaved by caspase-3 is located in the hinge region. (B) The GFP-PKCδ viral construct expresses PKCδ with EGFP fused to the C-terminus of the protein. (C) The CM-GFP-PKCδ viral construct expresses PKCδ with D327 in the caspase-3 cleavage site mutated to A327 and EGFP fused to the C-terminus.
